# Pleiotropic fitness effects of the lncRNA *Uhg4* in *Drosophila melanogaster*

**DOI:** 10.1101/2022.05.26.493587

**Authors:** Rebecca A. MacPherson, Vijay Shankar, Lakshmi T. Sunkara, Rachel C. Hannah, Marion R. Campbell, Robert R. H. Anholt, Trudy F. C. Mackay

## Abstract

Long noncoding RNAs (lncRNAs) are a diverse class of RNAs that are critical for gene regulation, DNA repair, and splicing, and have been implicated in development, stress response, and cancer. However, the functions of many lncRNAs remain unknown. In *Drosophila melanogaster*, *U snoRNA host gene 4 (Uhg4*) encodes an antisense long noncoding RNA that is host to seven small nucleolar RNAs (snoRNAs). *Uhg4* is expressed ubiquitously during development and in all adult tissues, with maximal expression in ovaries; however, it has no annotated function(s). Here, we used CRISPR-*Cas9* germline gene editing to generate multiple deletions spanning the promoter region and first exon of *Uhg4*. Females showed arrested egg development and both males and females were sterile. In addition, *Uhg4* deletion mutants showed delayed development and decreased viability, and changes in sleep and responses to stress. Whole-genome RNA sequencing of *Uhg4* deletion flies and their controls identified co-regulated genes and genetic interaction networks associated with *Uhg4*. Gene ontology analyses highlighted a broad spectrum of biological processes, including regulation of transcription and translation, morphogenesis, and stress response. Thus, *Uhg4* is a lncRNA essential for reproduction with pleiotropic effects on multiple fitness traits.

## INTRODUCTION

Long noncoding RNAs (lncRNAs) are a diverse class of non-coding RNAs of at least 200 nucleotides in length. Although lncRNAs were initially thought to be “junk” due to their noncoding status, we now know that lncRNAs are critical for various biological processes, including, but not limited to, transcription and gene regulation [1–8], chromatin architecture [9–11], DNA damage response [12–14], and scaffolding and nuclear organization [15–18]. LncRNAs can also act as miRNA sponges [19–21], regulate gene splicing [17,22–25], and be translated into functional peptides [26–28]. LncRNAs can be localized in the nucleus, cytoplasm, or to a specific organelle [29–32], can act in *cis* or in *trans* [8,23,33], and may be conserved across taxa in sequence and/or function [34–36]. Across taxa, lncRNA dysregulation has been implicated in cancer [37–40], neurological disorders [6,41–43], and immune and stress response [44–47]. Additional studies in *Drosophila melanogaster* have demonstrated critical roles for lncRNAs in development [48–53], gonadal function [48,54–56], sleep [19], locomotion [57], and courtship behavior [58, 59].

Roles for lncRNAs in development, viability, and fertility have been identified in multiple model systems [4,8,34,55,60,61]. However, fitness roles for some mammalian lncRNAs, including lncRNAs previously linked to fitness traits with gene-knockdown experiments, have not been replicated using CRISPR-based cell line or animal model knockouts [62–65]. Other recently-discovered mammalian lncRNAs with expression limited to reproductive tissues do not affect reproductive phenotypes in knockout mice [66, 67]. This controversy also extends beyond mammalian systems. Drosophila and zebrafish studies on CRISPR-mediated deletions of developmentally-expressed lncRNAs – some of which were previously implicated in development via RNA interference or morpholino-induced knockdown studies – also failed to identify roles for these lncRNAs in development, viability, or embryogenesis after no overt phenotypes were detected in knockout animals [68–70]. Inconsistencies across lncRNA studies may be due to the method of gene perturbation (*e.g.,* RNAi interference, CRISPR-*Cas9*), discordant phenotyping, transcriptional noise competing with low expression of some lncRNAs, differences in the genetic background of organisms used across studies on the same lncRNA, and/or functional redundancy of the lncRNA [61,69,71,72]. Furthermore, given the abundance of sex-specific and tissue-specific lncRNA expression, it is also possible that broader developmental pathways may overshadow effects due to loss of a lncRNA or be limited to a single tissue or behavioral phenotype [61, 72].

Here, we evaluate the effects of loss of function alleles of the *D. melanogaster* gene encoding *U snoRNA host gene 4* (*Uhg4*; FBgn0083124). *Uhg4* is a lncRNA with unknown function and is the host gene for seven intronic small nucleolar RNAs (snoRNAs), which guide posttranscriptional ribosomal RNA modification and processing [73, 74]. Some *U snoRNA* host genes interact with regulatory proteins controlling *piwi*-interacting RNAs (piRNAs) [75], which are noncoding RNAs involved in transposon silencing in germ cells [76, 77]. Although adult expression of *Uhg4* is highest in ovaries, it is expressed ubiquitously during development and in other adult tissues, including the brain and central nervous system, fat body, and trachea [78, 79]. *Uhg4* expression is correlated with modulation of expression of a subset of snoRNAs in response to developmental alcohol exposure in Drosophila females [80]. We used CRISPR/*Cas9* germline gene editing to create deletions in the promoter region and first exon of *Uhg4.* These mutations have pleiotropic effects on fitness-related traits, stress responses, sleep and activity phenotypes, and transcript abundances of both non-coding and protein-coding RNAs.

## RESULTS

### Generation of *Uhg4* Null Alleles

*Uhg4* is a long noncoding RNA that is host to seven small nucleolar RNAs (snoRNAs), and is expressed ubiquitously throughout development and in adults, with the highest adult expression in ovaries [78, 79]. We used CRISPR*-Cas9* and a double guide RNA vector to target the deletion of a ∼685bp region that includes the promoter region upstream of *Uhg4* as well as the first exon of *Uhg4* (Figure 1) in two DGRP lines, DGRP_208 and DGRP_705. We isolated seven independent deletion mutations in DGRP_208, most of which varied from one another by a few base pairs (Figure 1). The DGRP_208 deletions spanned the promoter region and the first exon of *Uhg4*. We also isolated four independent identical mutations in DGRP_705. The DGRP_705 deletions removed the *Uhg4* promotor region and retained the first exon, and included a 44bp AT-rich insertion in the first *Uhg4* intron upstream of the start of *snoRNA:Psi28S-2949* (Figure 1). Sanger sequencing shows that the small nucleolar RNA (snoRNA) *snoRNA:Psi28S-2949* within the first exon of *Uhg4* is intact for all independently obtained deletions (Figure 1).

**Figure 1.**
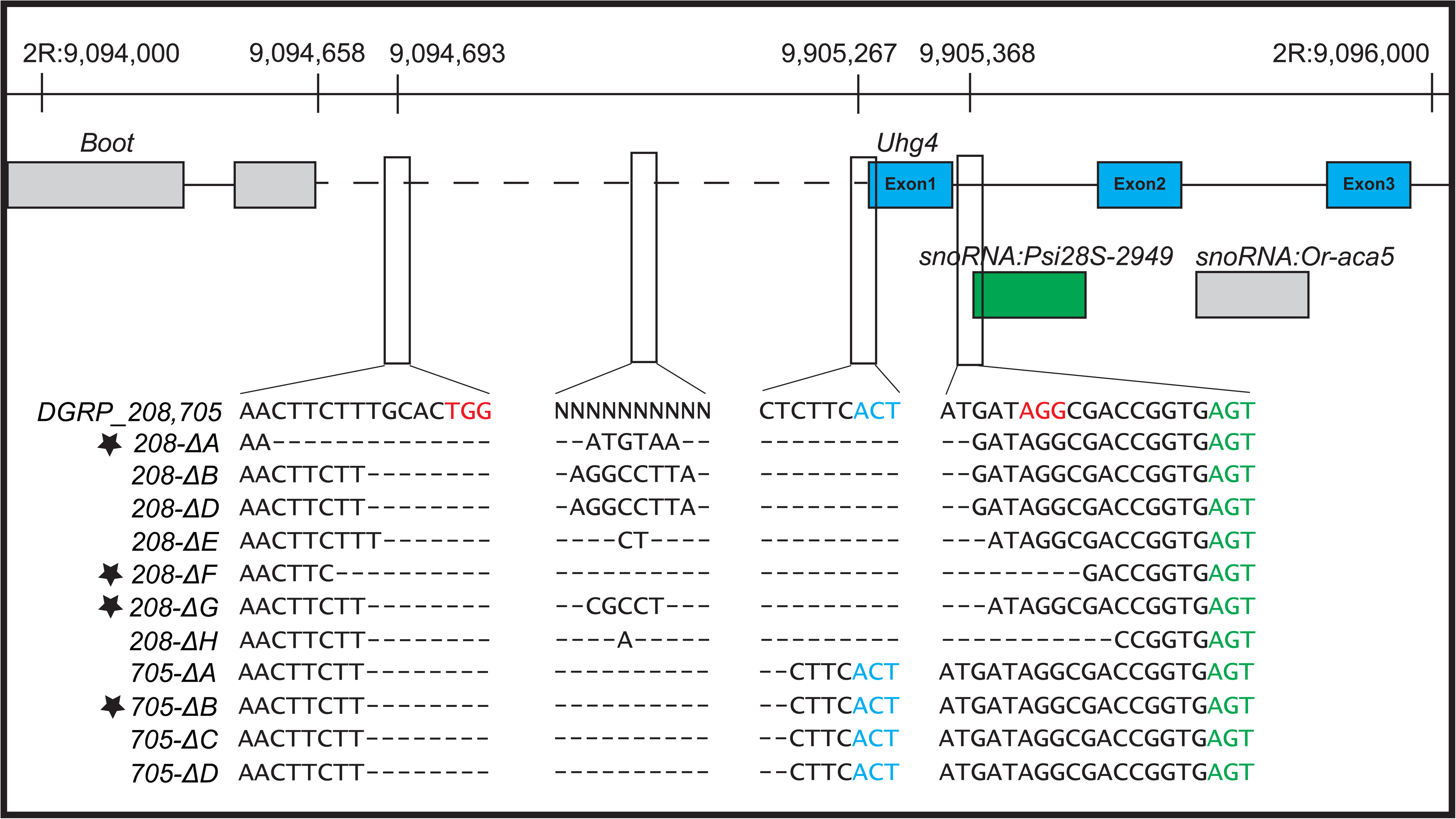
Uhg4 deletions. Diagram showing deletions across the *Uhg4* locus with Sanger sequencing data for the wildtype (DGRP_208, DGRP_705) and mutant genotypes. Genomic coordinates are shown at the top of the figure. The blue and green font colors indicate nucleotides of *Uhg4* and *snoRNA:Psi28S*-2949 coding regions, respectively. Red font colors indicate the PAM sites. “-“ indicates deleted nucleotides and N refers to multiple different nucleotides between the two wild-type DGRP lines. Genotypes used for phenotypic analyses are designated with a star.

Here, we focus on deletions *Uhg4^208-ΔA^*, *Uhg4^208-ΔF^*, *Uhg4^208-ΔG^*, and *Uhg4^705-ΔB^*, hereafter referred to as *208-ΔA*, *208-ΔF*, *208-ΔG,* and *705-ΔB*, respectively. We randomly selected these mutations for further study after several generations of backcrossing to the original genetic background. Both genetic backgrounds are highly inbred and free of inversions, and DGRP_208 is free of *Wolbachia* infection [81]. Flies homozygous for the DGRP_208 *Uhg4* deletions show changes in their resting wing position (File S1, S2). Compared to the control, *Uhg4* deletion flies carry their wings in an elevated and partially horizontally spread position.

### Effects of *Uhg4* Deletions on Fitness Traits

#### Fertility and Mating Behavior

When we generated the *Uhg4* deletion fly stocks, we did not observe eggs in vials that contained only flies homozygous for a *Uhg4* deletion and we needed to use a *CyO* balancer chromosome to maintain the deletion lines (Figure S1). To test whether the lack of eggs could be due to a failure of the *Uhg4* deletion flies to mate, we assessed mating latency and copulation duration and found that, although deletion flies do not produce progeny when mated, they do exhibit normal mating latencies, copulation times, and proportion of flies mated (Figure S2, Tables S1A, S1B). After crossing flies with the deletion to wild-type DGRP_208 flies of the opposite sex, we observed that *Uhg4* deletion females do not lay eggs, regardless of mating status or genotype of the male partner, and wild-type eggs fertilized by *Uhg4* deletion males do not develop past the embryo stage. To further probe why *Uhg4* deletion females are sterile, we dissected their ovaries after mating. We did not observe any late-stage (> 11) oocytes in ovaries of *Uhg4* deletion females, whereas ovaries in control DGRP_208 flies do contain late-stage oocytes with dorsal filaments (Figure 2). We did not perform these analyses for the *705-ΔB Uhg4* allele because it is nearly lethal as a homozygote, with rare viable adults. However, the few escaper *705-ΔB* females did not lay eggs. These results show that *Uhg4* is critical for fertility in both sexes and that the sterility of *Uhg4* deletion in females may stem from a lack of fully developed eggs.

**Figure 2.**
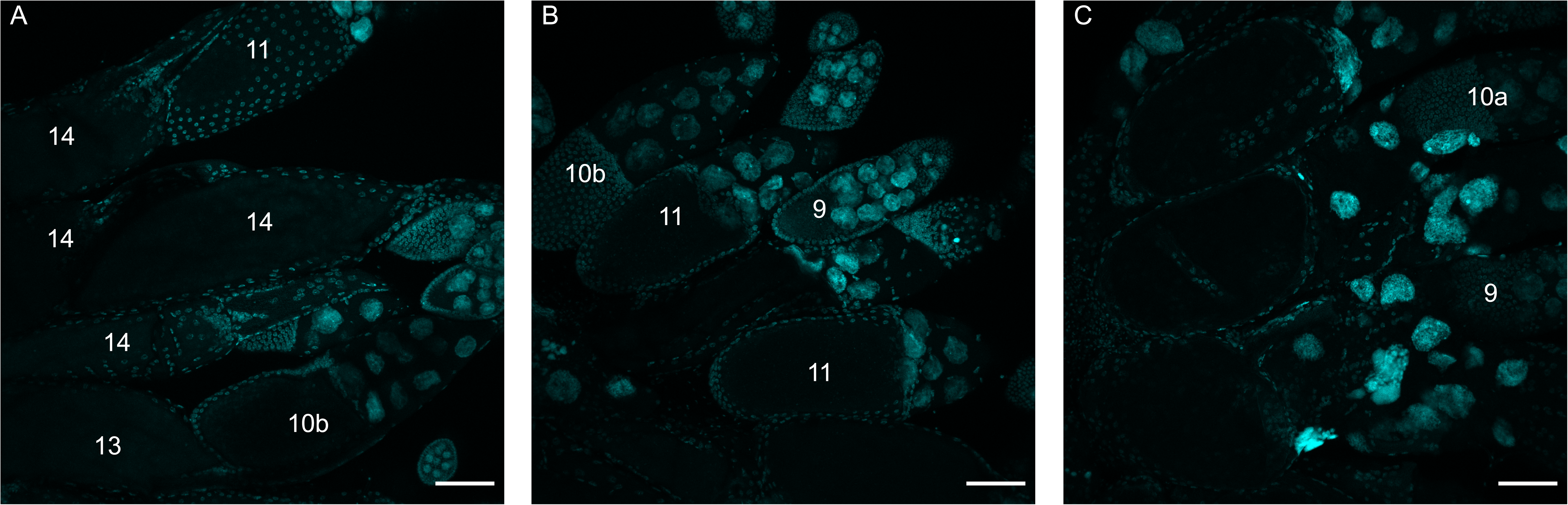
Late-stage oocytes are absent in *Uhg4* deletion lines. Maximum projection z-stack images (60 slices) of DAPI-stained developing egg chambers. (A) Control *Uhg4* (DGRP_208) ovaries. (B) *208-ΔF* ovaries. (C) *208-ΔG* ovaries. Compared to wild type, late-stage oocytes (stage number > 11) and any dorsal filaments are absent in the ovaries of flies with a *Uhg4* deletion. Scale bars represent 75µm. Numbers represent stages based on Jia et al. [110].

#### Development Time and Viability

During maintenance of *CyO*/*Uhg4-*deletion flies, we observed delayed emergence and skewed non-Mendelian ratios of *CyO*/deletion heterozygotes and homozygous deletion progeny. We formally assessed egg-adult development time and viability for the DGRP_208 *Uhg4* deletion lines compared to the wild-type control. We placed 50 eggs from matings of *208-ΔA*/*CyO*, *208-ΔF*/*CyO*, *208-ΔG*/*CyO* or DGRP_208/*CyO* (wild type control) flies in 25 vials each and recorded the day each fly emerged, as well as the sex, balancer genotype (*Cy* or straight wing), and the total number of flies for each sex and balancer genotype that emerged. Compared to the wild-type controls, flies with *Uhg4* deletions have delayed development by about one day (*p* < 0.0001, Figure 3A, Table S1C). Furthermore, *208-ΔA* males and all *Uhg4* deletion females show a 2.0- to 5.7-fold decrease in viability (*p* < 0.0001 for all lines, Figure 3B, Table S1D, S1E). These results suggest that *Uhg4* is necessary for the normal timing of egg-adult development and viability.

**Figure 3.**
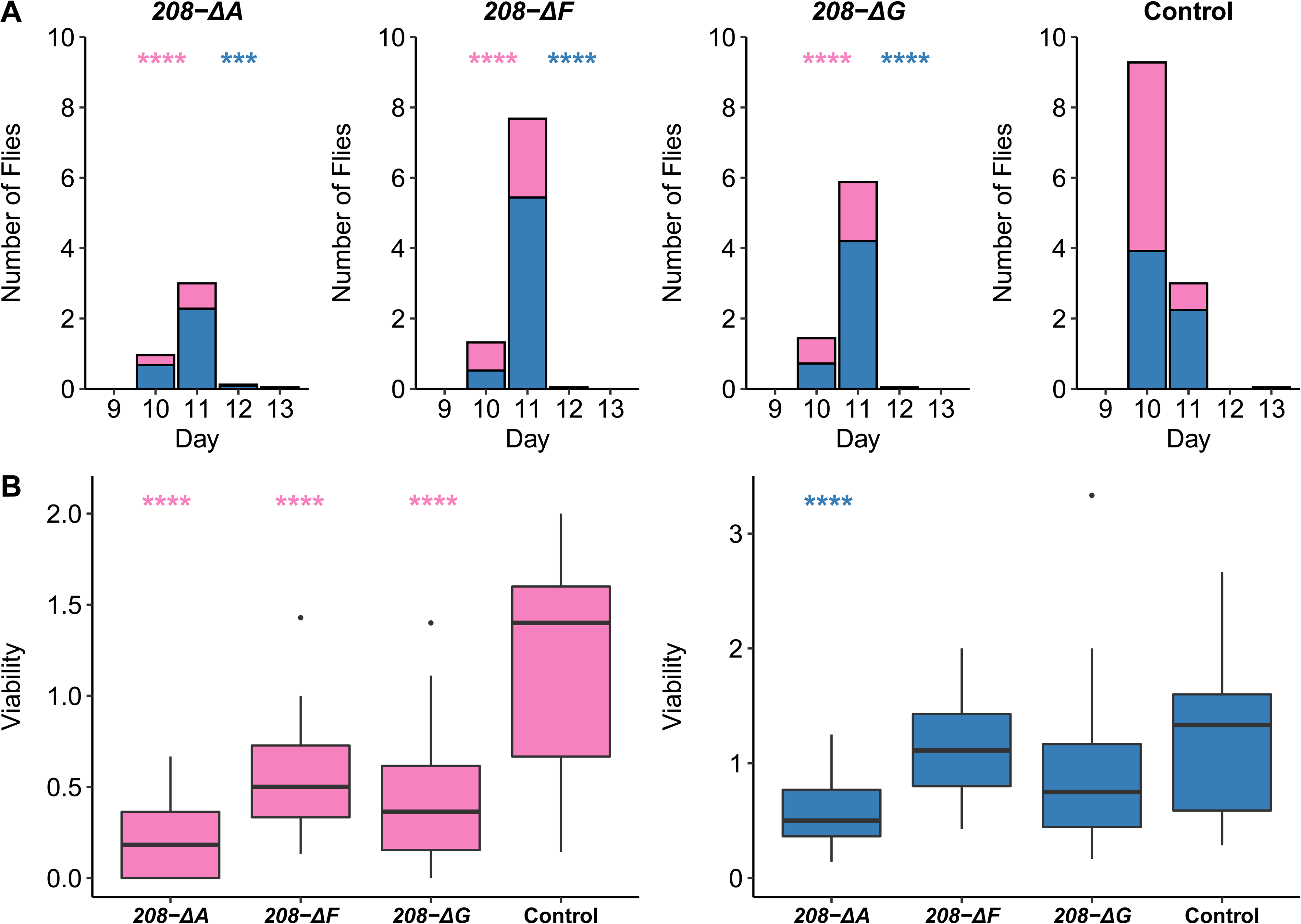
Effects of *Uhg4* deletion on egg-adult development and viability. (A) Stacked bar plots showing the average number of flies homozygous for *208-ΔA*, *208-ΔF*, *208-ΔG*, or wild type *Uhg4* (DGRP_208) that emerge from crosses of *208-ΔA*/*CyO*, *208-ΔF*/*CyO*, *208-ΔG*/*CyO* and DGRP_208/*CyO* flies. (B-C) Boxplots displaying viability coefficients for females (B) and males (C) for *Uhg4* deletion lines (*208-ΔA*, *208-ΔF*, *208-ΔG*) and the wild type (DGRP_208). Males are shown in blue and females are shown in pink. *N* = 25 vials of 50 embryos each per genotype. See Table S1 for all ANOVAs. *p*-values on the figure are for the comparisons of each sex to the control, by genotype. *** *p* < 0.001, **** *p* < 0.0001

### Effects of *Uhg4* Deletions on Responses to Stressors and Sleep Traits

#### Stress Responses

We assessed the effect of several stress conditions (heat shock, chill coma recovery, and ethanol sedation) on *Uhg4* deletion and wild-type control flies (Figure 4). On average, *Uhg4* deletion lines take longer to recover from a chill-induced coma than the wild-type in analyses pooled across sexes and all deletion genotypes (*p* = 0.013, Table S1F), although there is no difference in the proportion of flies that recover from a chill-induced coma for the deletion lines compared to the control (Table S1G). However, the response to a chill-induced coma is both sex- and genotype-specific. The chill coma recovery time for all deletion lines compared to the wild type is significant for males (*p* = 0.0015) but not females (*p* = 0.666); and males are only significant for the *208-ΔA* (*p* < 0.0001) and *208-ΔF* (*p* = 0.0007) deletion genotypes (Figure 4A). The *Uhg4* deletion lines also have reduced survival on average following a heat shock than the wild type in analyses pooled across sexes and all deletion genotypes (*p* = 0.0016, Table S1F). These effects were genotype-specific as well; only *208-ΔF* (*p* = 0.0013) and *208-ΔG* (*p* = 0.034) were formally significantly different from the wild type, although *208-ΔA* trended in the same direction (*p* = 0.093) (Figure 4B). In contrast to temperature-related stressors, *Uhg4* deletion flies show decreased susceptibility to ethanol-induced stress. The time to sedation in response to acute ethanol exposure is significantly increased averaged over all *Uhg4* deletion lines compared to the wild type in the analyses pooled across sexes (*p* < 0.0001) as well as in females (*p* = 0.0007) and males (*p* < 0.0001) (Table S1F). Although all *Uhg4* deletion sex/genotype comparisons had significantly increased sedation times relative to the control, there was heterogeneity in the magnitudes of effects among the deletion genotypes and sexes. In females, *208-ΔF* (*p* < 0.0001) had larger effects than *208-ΔA* (*p* = 0.016) and *208-ΔG* (*p* = 0.036); while in males all *Uhg4* deletion lines had similar effects (*p* < 0.0001 for all) (Figure 4C).

**Figure 4.**
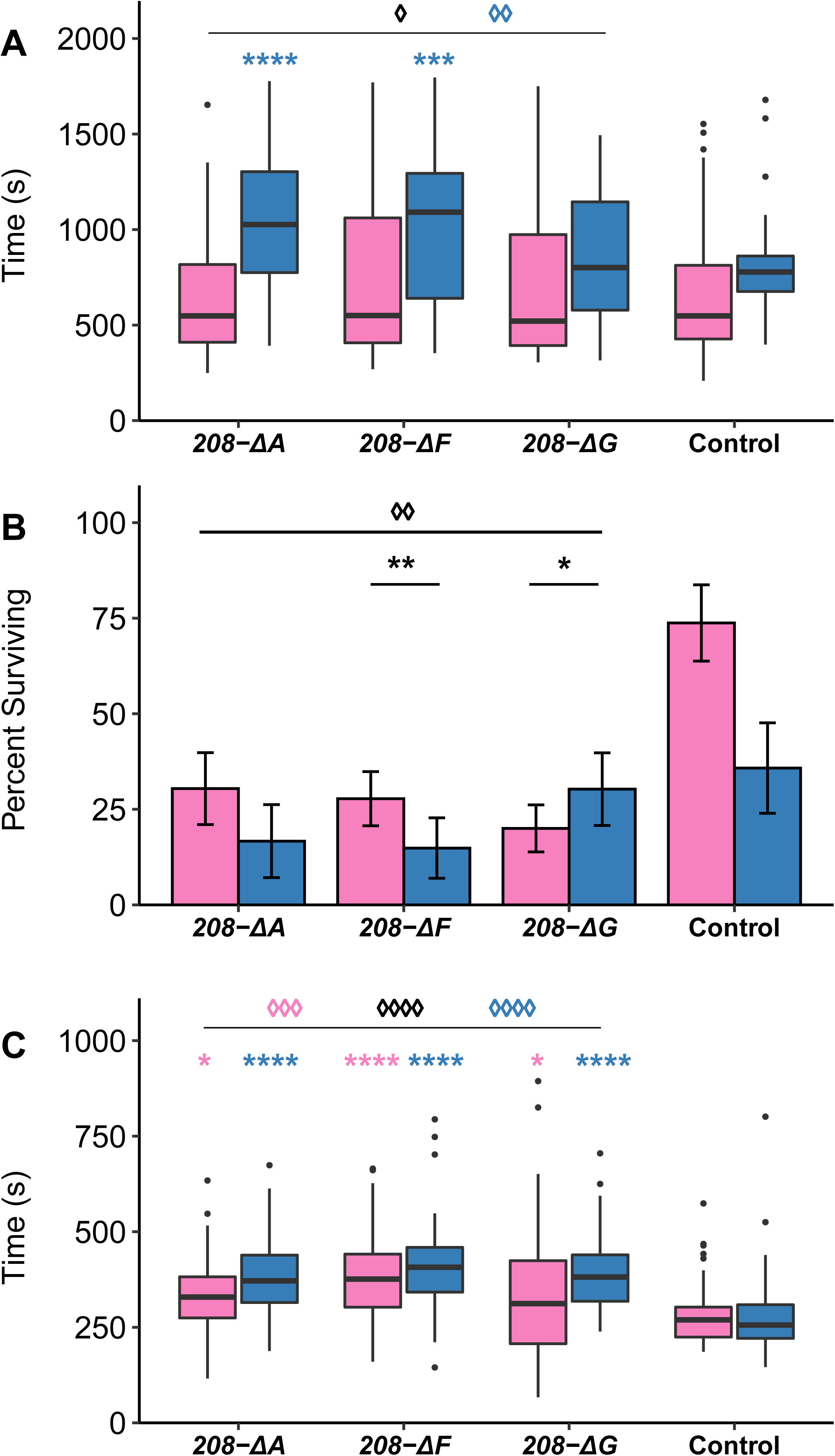
Effects of *Uhg4* deletion on responses to stress. (A) Boxplots indicating chill coma recovery time (n = 48-57 flies per sex and genotype). (B) Bar plots showing the average proportion of flies surviving heat shock (n = 10 replicates of 9 flies per sex and genotype). (C) Boxplots showing ethanol sedation sensitivity time (n = 44-52 flies per sex and genotype). Blue boxes indicate males and pink boxes indicate females. Error bars indicate standard error. See Table S1 for all ANOVAs. *p*-values on the figure indicated by an asterisk (*) represent comparisons of each sex to the control, by genotype. *p*-values on the figure indicated by a diamond (◊) represent comparisons of pooled *Uhg4* deletion genotypes versus the control. * *p* < 0.05, ** *p* < 0.01, *** *p* < 0.001, **** *p* < 0.0001

#### Sleep and Activity

We used the Drosophila Activity Monitor System to assess the effects of *Uhg4* deletions on sleep and activity traits. *208-ΔA*, *208-ΔF*, and *208-ΔG* flies sleep more during the day (*p* = 0.0009 from the analysis of all deletion lines pooled across sexes compared to the wild type, Figure 5A, Table S1H) and night (*p* = 0.031 from the analysis of all deletion lines pooled across sexes compared to the wild type, Figure 5B, Table S1H). The effects of the deletions are much greater on day than on night sleep. In addition, day sleep is significant averaged over all deletion lines for males (*p* = 0.026) and females (*p* = 0.014); while the effect on night sleep is male-specific (*p* = 0.011) (Table S1H). Although the *Uhg4* deletions sleep longer than the wild type during the day and night, they also have more fragmented sleep, as the number of sleep bouts increases both during the day (*p* = 0.002 from the analysis of all deletion lines pooled across sexes compared to the wild type, Figure 5C, Table S1H) and night (*p* = 0.0001 from the analysis of all deletion lines pooled across sexes compared to the wild type, Figure 5D, Table S1H). Concomitant with increased day and night sleep, the *Uhg4* deletions on average have decreased length of activity bouts during the day (*p* = 0.0061, Figure 5E, Table S1H as well as decreased total locomotor activity (*p* = 0.0062 from the analysis of all deletion lines pooled across sexes compared to the wild type, Figure 5F, Table S1H).

**Figure 5.**
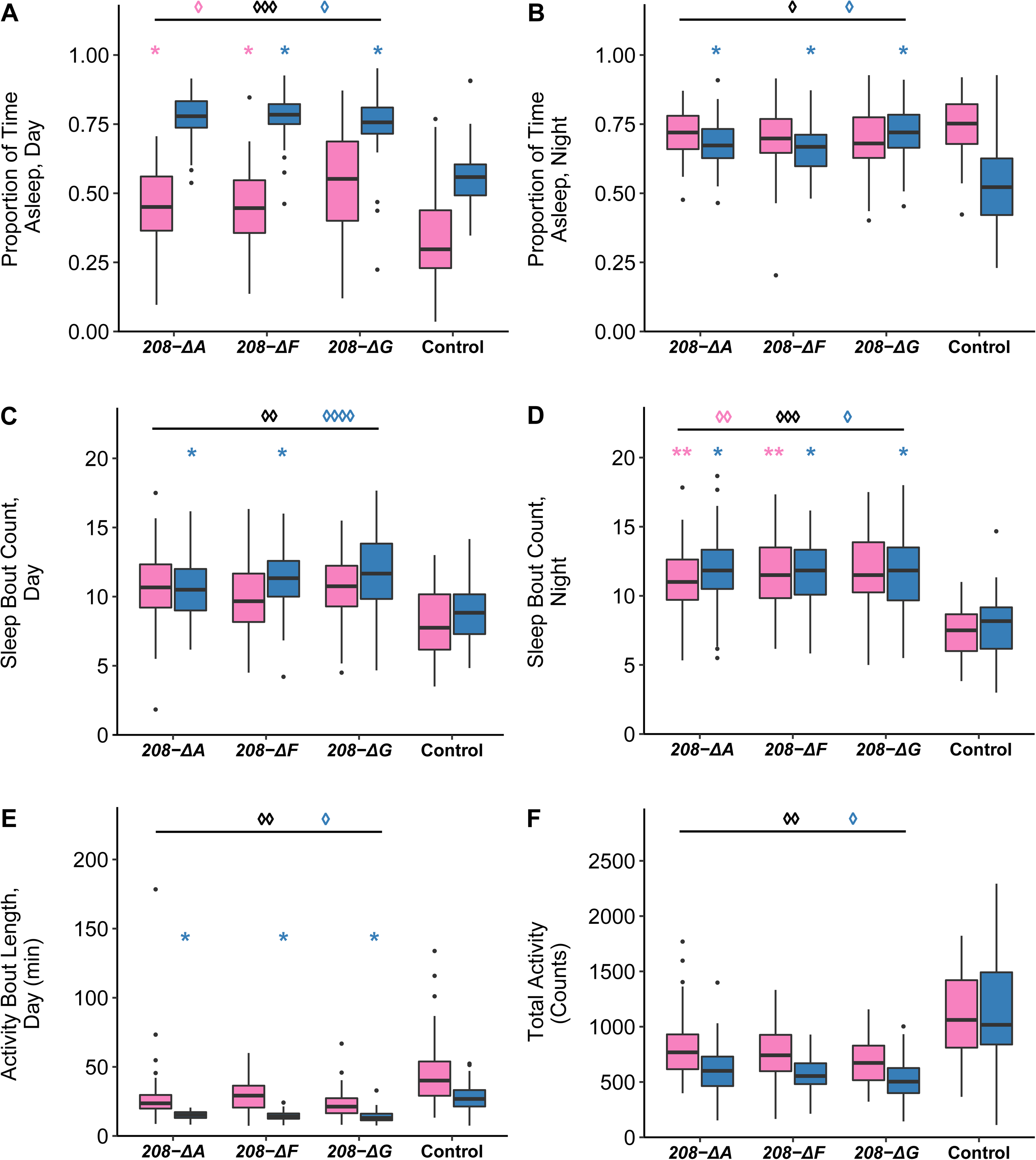
Effects of *Uhg4* deletion on sleep and activity phenotypes. (A) Boxplots showing the proportion of daytime sleep, (B) the proportion of nighttime sleep, (C) the number of sleep bouts at night, (D) the number of sleep bouts during the day, (E) total activity, and (F) activity bout length during the day. Blue boxes indicate males and pink boxes indicate females. Day hours are from 6 am-6 pm. N = 61-64 flies per sex per line, except *208-ΔG*, which had only 32 females. See Table S1 for ANOVAs. *p*-values on the figure indicated by an asterisk (*) represent comparisons of each sex to the control, by genotype. *p*-values on the figure indicated by a diamond (◊) represent comparisons of pooled *Uhg4* deletion genotypes versus the control. * *p* < 0.05, ** *p* < 0.01, *** *p* < 0.001, **** *p* < 0.0001

**Figure 6.**
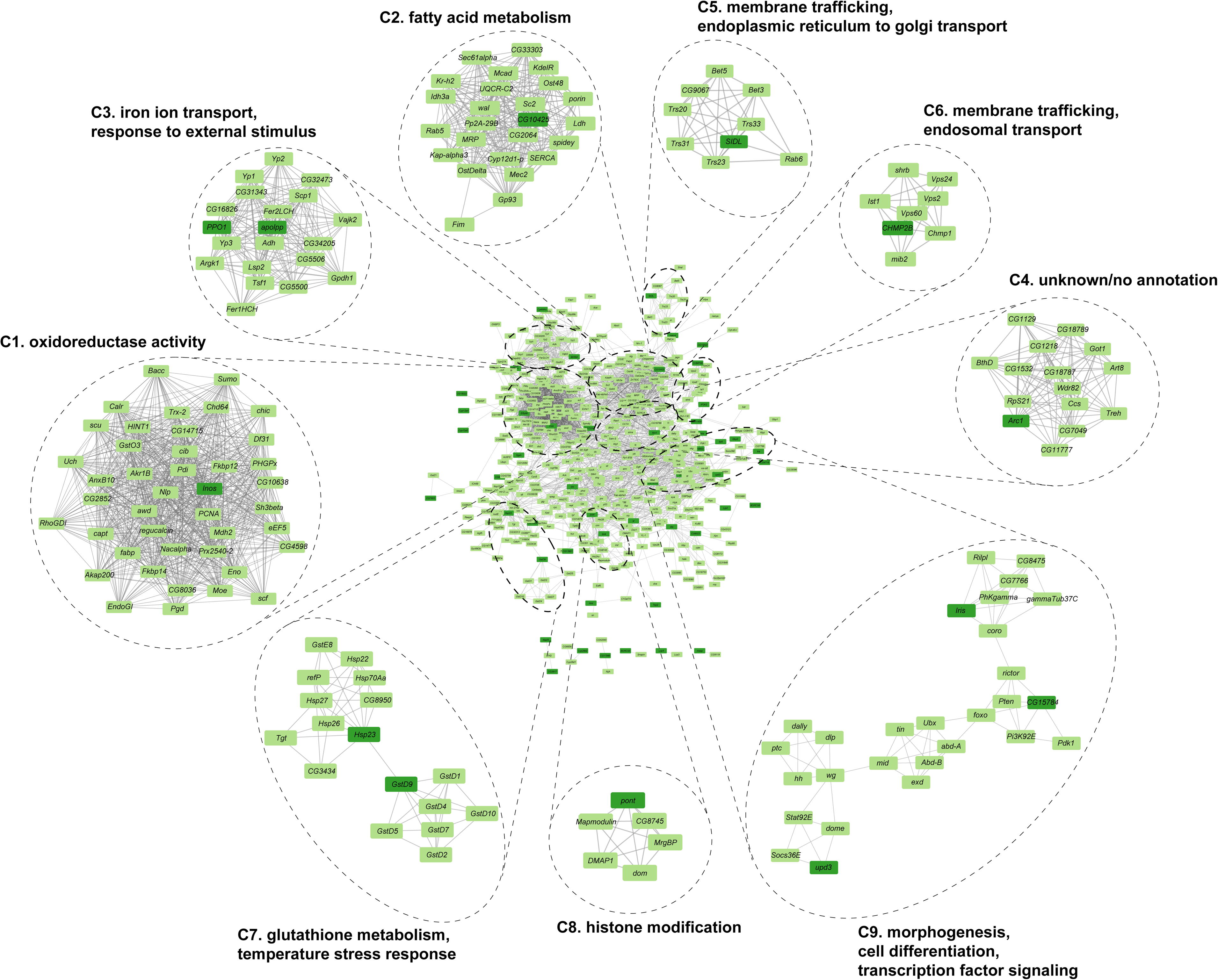
Interaction networks of genes coregulated with *Uhg4*. Interaction networks (physical and genetic interactions) based on 180 genes with differential expression for the *Line* term (BH-FDR<.1). The full network (including interaction neighbors within 1 degree) is in the middle of the figure, with MCODE subclusters and subcluster GO enrichment annotations (Table S4) on the perimeter. Dark green indicates the input 180 genes, and light green indicates an interaction neighbor. Names are Drosophila gene symbols. Numbers in parentheses indicate the cluster number.

### Effects of *Uhg4* Deletions on Genome-Wide Gene Expression

To assess the consequences of deletion of *Uhg4* on genome-wide gene expression, we performed RNA-sequencing on *208-ΔA*, *208-ΔF*, *208-ΔG,* and DGRP_208 whole flies, separately for males and females. We performed factorial fixed effect ANOVAs for each of the 16,212 genes expressed in young adult flies that evaluate the significance of the main effects of the four *Uhg4* genotypes (Line), Sex, and the Line by Sex interaction. Plotting ordered raw *p*-values and BH-FDR adjusted *p*-values against the number of tests revealed a non-monotonic relationship between raw *p*-values and adjusted *p*-values (Figure S3a). This relationship caused an artificial inflation in the number of differentially expressed genes at BH-FDR < 0.1. Therefore, we used a BH-FDR thresholding approach to identify statistically significant genes at BH-FDR < 0.1. Briefly, after ordering the genes based on ascending raw *p*-values, we compared each gene’s raw *p*-value to its BH-FDR critical value calculated as 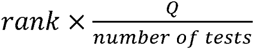 at both Q = 0.05 and Q = 0.1 (Figure S3b). For both critical values, *p*-value thresholds were determined as the first occurrence of the raw *p*-value greater than critical values. Using this method, we identified 17 differentially expressed genes at a BH-FDR < 0.05 for the Line and/or Line by Sex terms (Table S2, Table S3). The top three differentially expressed genes were *Uhg4*, *snoRNA:Psi28S-2949,* and *snoRNA: Or-aca5*. The near-complete loss of *Uhg4* expression in the deletion genotypes is expected due to the deletions of the promoter region and exon 1. The two snoRNAs are located in the first two introns of *Uhg4*. Decreased expression of *snoRNA:Psi28S-2949* and *snoRNA:Or-aca5* suggests the *Uhg4* deletions affect regulatory sequences common to both snoRNAs, since their coding sequences are not altered (Figure 1). Ten of the 17 differentially expressed genes are computationally predicted genes and/or noncoding RNAs, including *Uhg4,* and have limited to no information on gene function. One of the 17 significantly differentially expressed genes was *insulin-like peptide 6* (*Ilp6*).

A total of 17 genes that are differentially expressed in *Uhg4* null genotypes is not sufficient to infer the function of *Uhg4* from the enrichment of Gene Ontologies (GO) and networks of the co-regulated genes. Therefore we relaxed the significance threshold to BH-FDR < 0.1 for the Line and Line by Sex terms of the ANOVA models. This resulted in 180 differentially expressed genes (Table S2). Notably, all genes significant for the Line by Sex terms also had significant Line effects (Table S3). For these 180 genes, only one GO term (humoral immune response GO:0006959) was enriched at BH-FDR < 0.05 (Table S4). However, 47 of the 180 genes did not map to a GO term. These analyses suggest that although deletion of *Uhg4* does change the transcriptome, many of these changes are associated with genes about which little, if anything, is known.

To fully quantify transcriptomic changes in the *Uhg4* deletion flies, we also assessed changes in expression of novel transcribed regions (NTRs) previously identified in the DGRP (including DGRP_208) [82]. Of the 18,581 NTRs analyzed, we found three (12) were differentially expressed at BH-FDR < 0.05 (0.1) for the Line term, and one NTR was differentially expressed for the Line by Sex term at BH-FDR < 0.1. Together with the protein coding genes, we have a total 193 differentially expressed genes/NTRs in the *Uhg4* deletion lines.

We then performed k-means clustering to assess patterns of expression across genotypes, using k = 8, as it offered the largest number of clusters without redundancy of expression patterns across clusters. GO analysis of these k-means clusters emphasizes the sparsity of information currently known about the genes that are coregulated with *Uhg4*. Of the eight clusters, only four clusters (clusters 1, 3, 6, 7) have enriched GO terms (BH-FDR < 0.05). Cluster 1 is enriched for genes involved in immune response, cluster 3 is enriched for genes involved in response to stress and temperature, and clusters 6 and 7 are enriched for genes involved in cuticle structure (Table S4). Cluster 1 consists of genes largely downregulated in deletion lines compared to the control, including *Uhg4*, as well as other noncoding RNAs and two NTRs (Figure S4). Broadly, cluster 2 contains sexually dimorphic genes, whereas clusters 3 and 5 show genes upregulated in *208-ΔA* and *208-ΔG*, respectively, compared to other lines. *208-ΔF* displays intermediate expression patterns compared to *208-ΔA* and *208-ΔG* in clusters 3, 6, and 8 (Figure S4).

In an effort to generate further hypotheses about the role of *Uhg4* and its co-regulated genes, we used the FlyBase database of known genetic and physical interactions within *D. melanogaster* [79] to generate interaction networks for the subset of 180 co-regulated genes at BH-FDR < 0.1. We also included first-degree interaction neighbors, genes, or proteins that are recorded in the database as directly interacting with at least one of the 180 focal genes. These networks revealed nine subclusters containing genes enriched for a broad spectrum of biological processes, including iron ion transport, fatty acid metabolism, temperature stress response, membrane trafficking, and morphogenesis (Figure 4, Table S4). Cluster 9, which contains the genes *Ubx*, *dlp*, and *Pten*, among others, was enriched for hundreds of GO terms, far more than any other cluster, suggesting genes in this cluster are critical for a wide range of processes, including morphogenesis, cell differentiation, transcription factor signaling, sleep and activity, reproduction, stress response, and metabolism (Table S4). Cluster 4 was not enriched for any GO terms and attempts at manual annotation did not reveal related functions for genes in this cluster (Table S4). These results indicate that the lncRNA *Uhg4* contributes to diverse cellular functions.

## DISCUSSION

We used *Uhg4*-knockout flies to assess the role of the lncRNA *Uhg4* across multiple fitness traits. We present evidence that *Uhg4* is critical for egg development and fertility and has pleiotropic effects on viability, development, stress responses, and sleep. Genome-wide gene expression data support the finding that *Uhg4* has pleiotropic effects, as *Uhg4* is coregulated with genes involved in a wide range of biological processes, including development, trafficking, metabolism, and stress response. We also observed that many of the genes differentially expressed upon loss of *Uhg4* are also noncoding RNAs in addition to predicted genes, emphasizing the need for further study of the effects on fitness traits of noncoding RNAs in *D. melanogaster*.

Although many of the coregulated genes themselves do not have associated GO terms, many biological processes and functions implicated by GO analysis align with observed changes in organismal phenotypes. Terms involving response to external stimuli such as temperature and ethanol, as well as terms relating to immune system response, are enriched in multiple k-means and interaction network clusters. These terms align with the increased susceptibility of the *Uhg4* deletion flies to extreme hot and cold temperatures, as well as the decreased susceptibility to ethanol-mediated stressors. Interestingly, *Alcohol dehydrogenase* (*Adh*) interacts with at least two genes coregulated with *Uhg4­,* providing a possible mechanistic link between *Uhg4* and ethanol response; *Uhg4* has been previously implicated in developmental ethanol exposure, which also results in decreased susceptibility to ethanol exposure in adult flies [80]. Although not annotated in Figure 4, interaction cluster nine is enriched for genes involved in wing morphogenesis and gamete generation (Table S4), which could explain the changes to wing position and sterility phenotypes, respectively, observed in *Uhg4* deletion flies. Cluster 9, and its more stringent 2-degree interaction counterpart (Cluster 3 in Figure S5), are enriched for hundreds of GO terms (Table S4), indicating that these genes have wide-ranging impacts. Thus, the effects of *Uhg4* deletion may extend to additional traits that we did not assess, such as iron ion transport, or other intracellular phenotypes.

Based on the DAPI-stained ovary images (Figure 2) showing a lack of late-stage oocytes, we hypothesize that *Uhg4*-deletion females are capable of laying eggs but do not develop late-stage oocytes. This oocyte development phenotype is supported by the high expression of *Uhg4* in ovaries. However, this does not explain why embryos from wild-type females and *Uhg4* deletion males fail to develop.

*Bootlegger* (*Boot*), a gene located immediately upstream of and in opposite orientation to *Uhg4*, is also critical for proper egg development [83, 84]. Our Sanger sequencing data indicate that *Boot* is intact in all deletions and our RNAseq data do not show *Boot* to be differentially expressed. Furthermore, unlike *Uhg4*, *Boot* is minimally expressed in males and would unlikely be responsible for the sterility observed in *Uhg4*-deficient males. We are therefore confident that the phenotypes observed in our *Uhg4* deletion flies can be attributed to *Uhg4*, though it is possible that *Uhg4* and *Boot* share promoter elements in their intergenic region.

The role of some snoRNA host genes such as *Uhg4* was thought to facilitate transcription and splicing of snoRNAs [85–87]. Based on our results, we hypothesize that *Uhg4* has roles independent of hosting snoRNAs, as most of its snoRNAs are not differentially expressed in *Uhg4* deletion flies. The abundance of genes/NTRs coregulated with *Uhg4* that are noncoding RNAs and/or have no known function makes speculation about the possible functional mechanisms by which *Uhg4* affects a wide range of pleiotropic phenotypes challenging. *Uhg4* could bind directly to DNA or transcription factors to modulate transcriptional regulation, acting in *cis* to regulate the expression of *snoRNA:Psi28S-2949* and *snoRNA:Or-aca5*, or in *trans* to regulate the expression of other genes we observed to be differentially expressed. *Uhg4* may also serve an oncogenic role, as overexpression of *Uhg4* in a Drosophila cell line is associated with tumor growth, and*Uhg4* is a downstream target of *Myc* [88]. *Uhg4* may also act to regulate gene expression at an epigenetic level via histone modifications (Cluster 8, Figure 4). Other lncRNAs in Drosophila that are important for thermotolerance are essential for remobilization of heterogenous ribonucleoprotein particles (hnRNPs) [41, 44]. *Uhg4*, as a host gene for snoRNAs, which also form ribonucleoproteins, might also modulate response to thermal stressors via ribonucleoproteins. *Uhg4* could also act in a similar manner to the transcript of *oskar*, which is critical for oogenesis in Drosophila [54,89,90]. *oskar* RNA facilitates oogenesis through multiple mechanisms, as it binds a translational regulator at one locus, has a separate 3’ region critical for proper egg-laying, and may also be involved in scaffolding of ribonucleoproteins [89].

In summary, our study shows that *Uhg4* is a lncRNA with pleiotropic functions and is indispensable for viability and reproduction in *D. melanogaster*.

## METHODS

### Generation of *Uhg4* Null Alleles

We used the flyCRISPR target finder [91] to design gRNAs flanking *Uhg4* Exon 1 (upstream: 5’-GAAGTAAAACTTCTTTGCACTGG -3’; downstream: 5’-GTAAGTATTATAGATATGATAGG -3’) that did not overlap with known genes and did not have predicted off-target effects, resulting in a 685bp deletion. We then synthesized a single gRNA vector containing both gRNAs [92]. Briefly, synthesized phosphorylated gRNAs with complementary sequences were ligated to form double-stranded gRNAs with CTTC overhangs. One double-stranded gRNA was cloned into *pBFv-U6.2* (Addgene #138400), and the other double-stranded gRNA was cloned into *pBFv-U6.2B* (Addgene #138401). Using flanking *Eco*RI and *Not*I sites, the resulting *U6* promoter and gRNA within *pBFV-U6.2* were excised and ligated into the *U6.2B* vector, creating a double-gRNA vector. Sanger sequencing confirmed the proper insertion of gRNAs.

The completed dual gRNA vector and *pBFv-nosP-Cas9* (Addgene #138402) were purified and co-injected (BestGene Inc., Chino Hills, CA) into at least 300 embryos from two different *D. melanogaster* Genetic Reference Panel (DGRP) lines [81, 93], DGRP_208 and DGRP_705. These lines have minimal heterozygosity and are homosequential for the standard karyotype for all common inversions; DGRP_208 is also free of *Wolbachia* infection [81].

To preserve the inbred genetic background of the DGRP lines, we screened flies for the presence of a deletion by individually isolating DNA from clipped wings of virgin female and male flies [94]. Fly wings were covered with 10 µL of 400 µg/mL protease K in a 10mM Tris-Cl at pH 8.2, 1mM EDTA and 25mM NaCl buffer and incubated at 37°C for 2h followed by 95°C for 2 min. We used 2μL of the resulting DNA mixture in a PCR reaction with primers (Left: 5’-CTAGCACGGAACCCTGGAAAT -3’; Right: 5’-GCAGCGCCTAGTAATCACAGA -3’) according to ApexRedTaq (Genesee Scientific, El Cajon, CA) manufacturer instructions, with a 61°C annealing temperature for 30 seconds and 30 cycles. The deletion mutations *Uhg4^208-ΔA^*, *Uhg4^208-ΔF^*, *Uhg4^208-ΔG^*, and *Uhg4^705-ΔB^* (hereafter referred to as *208-ΔA*, *208-ΔF*, *208-ΔG* and *705-ΔB*, respectively) were isolated and placed over a *CyO* balancer chromosome in the appropriate genetic background. *Uhg4* wild-type lines (DGRP_208 and DGRP_705) are used as controls in this study.

### Drosophila Culture

Flies were reared at 25°C with 50% humidity on standard cornmeal-molasses-agar medium (Genesee Scientific, El Cajon, CA), supplemented with yeast. Flies were maintained at controlled density on a 12-hour light-dark cycle (lights on at 6 am). Unless otherwise indicated, all behavioral assays were performed on 3-5 day old homozygous flies from 8:30 am to 11:30 am.

### Sanger Sequencing

DNA from homozygous mutant flies was sequenced by the Sanger chain termination method using a BigDye Terminator Kit v3.1 (ThermoFisher Scientific, Waltham, MA), with the same primers used to identify deletions. Sequencing was performed on an Applied Biosystems 3730xl DNA Analyzer (ThermoFisher Scientific, Waltham, MA).

### Quantitative Real-Time PCR

Flies were flash-frozen for RNA extraction at 3-5 days of age. Each sample contained 30 whole flies and was homogenized using a Fisherbrand™ Bead Mill (ThermoFisher Scientific, Waltham MA). RNA was extracted using the Direct-zol RNA Miniprep kit (Zymo Research, Irvine, CA), resuspended in RNase-free water, and kept at -80°C until further use. cDNA was synthesized using the iScript™ Reverse Transcription Supermix for RT-qPCR (Bio-Rad Laboratories, Inc., Hercules, CA) according to the manufacturer’s instructions. Quantitative real-time PCR to detect expression was performed on three biological replicates and two technical replicates per sample (except *208-ΔA*, which had two biological replicates), using SYBR™ Green PCR Master Mix (ThermoFisher Scientific, Waltham, MA) and primers (*Uhg4-*Forward: 5’-TCGGTCTTTCGATTTGGATT -3’; *Uhg4-*Reverse: 5’-TGTGTTAGTGAGCCACGTTTG -3’, spanning exons 4-5 of *Uhg4*; GAPDH-Left: 5’-CTTCTTCAGCGACACCCATT -3’; GAPDH-Right: 5’-ACCGAACTCGTTGTCGTACC -3’) according to the manufacturer’s instructions. Percent knockdown was assessed using *ΔΔct* [95]. No amplification was observed in non-template negative controls.

### Viability and Development Time

We placed 75 male-female pairs of flies from each of the DGRP_208 *Uhg4* genotypes (DGRP_208*/CyO*; *208-ΔA*/*CyO, 208-ΔF*/*CyO, 208-ΔG*/*CyO*) in large embryo collection cages (Genesee Scientific, San Diego, CA), supplemented with fresh yeast paste and grape juice-agar plates containing 3% agar and 1.2% sucrose in a 25% Welch’s Grape Juice concentrate solution every 12 hours. After 36 hours, we placed 50 0-12 hour embryos in each of 25 vials per genotype. From days 9-13 after egg-laying, we collected adult flies once daily and recorded the sex, wing phenotype, and day of emergence for each fly, including flies that eclosed, but died in the food. We calculated relative egg-adult viability (*v*) for each vial as the proportion of homozygous *Uhg4* adults (1-*r*) relative to the proportion of *Uhg4*/*CyO* heterozygote adults (*r*) as *v* = 2(1-*r*)/*r* [96].

### Fertility

We assessed the fertility of homozygous males and females from the DGRP_208 *Uhg4* deletion lines by crossing virgin deletion mutant flies to virgin DGRP_208 flies, respectively, of the opposite sex. In addition, we also performed crosses of males and females within each *Uhg4* deletion line. We set up four vials each with four males and four virgin females for each genotype. For each cross, qualitative observations (presence/absence) were made for each stage of development (embryos, first instar larvae, second instar larvae, third instar larvae, pupae, adult flies). We did not perform a formal fertility assay with the *705-ΔB* deletion line due to the very low viability of *705-ΔB* homozygotes.

### Mating Behavior

For each DGRP_208 *Uhg4* genotype (*208-ΔA*, *208-ΔF*, *208-ΔG*, DGRP_208), we placed 22-24 pairs of virgin flies in separate mating chambers [97, 98] to acclimate overnight. Between 8 am and 10 am, fly pairs were united and video-recorded for 30 minutes. Copulation duration and mating latency were recorded in seconds for each fly pair. We only included flies that mated in the analyses of mating behaviors and recorded the number of pairs that did not mate during the 30-minute testing period. We selected *208-ΔG* as a representative *Uhg4* mutant line and assessed the mating behaviors of crosses of *208-ΔG* males to DGRP_208 virgin females and *208-ΔG* virgin females to DGRP_208 males.

### Survival After Heat Shock

Vials with 2ml of medium and 9 homozygous flies per sex and *Uhg4* genotype were placed in a 37°C incubator for 2 or 3 hours beginning at 1 pm, with five replicate vials per sex per genotype per exposure time. After the heat shock, flies were transferred to fresh vials with 2ml of medium and allowed to recover overnight. Twenty-four hours after the heat shock, the number of surviving flies in each vial was recorded.

### Chill Coma Recovery Time

For each genotype we placed four vials containing 15 flies without medium on ice for 3 hours, sexes separately, and allowed the flies to recover in a 6-well cell culture plate (five flies per well, two genotypes per plate) for 30 minutes. We recorded the time until each fly righted itself by standing up [99]. Only flies that recovered within 30 minutes of being removed from the ice were included in the analysis (n = 48-57 flies per sex per line), although we also recorded the number of flies that did not recover.

### Ethanol Sensitivity

We assessed the time to sedation in response to acute ethanol exposure [100] on 44-52 flies per sex per *Uhg4* genotype. Briefly, flies were aspirated into a 24-well cell culture plate and placed opposite a 24-well cell culture plate containing 100% ethanol, separated by a layer of fine screen mesh. We recorded the time to sedation, defined as the moment each fly loses postural control.

### Sleep and Locomotor Activity

We collected sleep and locomotor activity data using Drosophila Activity Monitors (DAM) (TriKinetics, Waltham, MA). Briefly, we placed 1-2 day-old flies into DAM monitor tubes containing 2% agar with 5% sucrose, with two DAM monitors per sex per *Uhg4* genotype, for a total of 64 flies per sex and genotype. Sleep and activity data were recorded on days 3-8 of the fly lifespan on a 12-hour light-dark cycle. Sleep was defined as at least 5 minutes of inactivity. Only flies that survived the entire testing period were included in analyses, resulting in 61-64 flies per sex per *Uhg4* genotype (except for *208-ΔG* females, for which there was only one replicate of 32 individuals). The DAM data were initially processed with Shiny-RDAM [101] and resulting raw sleep and activity output files were downloaded for further statistical analysis.

### Statistical Analyses

Unless otherwise indicated, all behavioral assays were analyzed using the “PROC MIXED” command (for a mixed-effects model) or “PROC GLM” command (for a pure fixed-effects model) in SAS v3.8 as a Type III Analysis of Variance (ANOVA). Where appropriate, flies were randomized to avoid positional or time-related effects.

Development time (day of eclosion) was analyzed according to the model *Y* = *μ* + *L* + *S* + *G* + *L*×*S* + *L*×*G* + *S*×*G* + *L*×*S*×*G* + *Rep(L)* + *S*×*Rep(L)* + *G*×*Rep(L)* + *S×G×Rep(L)* + *ε*. Heat shock (percent of flies that survived) and all sleep and activity phenotypes (e.g. locomotor activity, bout count, percent of time asleep) were analyzed according to the model *Y* = *μ* + *L* + *S* + *L*×*S* + *Rep(L*×*S)* + *ε*. Chill coma recovery time (time to recover, in seconds) and ethanol sensitivity (time to sedation, in seconds) were analyzed according to the model *Y* = *μ* + *L* + *S* + *L*×*S* + *ε.* Mating latency (in seconds), mating duration (in seconds), and viability (viability coefficient *v*) were assessed according to the ANOVA model *Y* = *μ* + *L* + *ε*. In these models, *Y* is the phenotype of interest, *μ* is the overall mean, *L* is the fixed effect of Line (DGRP_208, *208-ΔA*, *208-ΔF*, *208-ΔG*), *S* is the fixed effect of sex (male, female), *G* is the fixed effect of balancer genotype (*Uhg4*/*CyO* heterozygote, *Uhg4* homozygote), *Rep* is replicate nested within lines, and *ε* is the residual error. We performed pairwise comparisons of each *Uhg4* deletion line (*208-ΔA*, *208-ΔF*, or *208-ΔG*) with the *Uhg4* wild type control, as well as a pooled comparison across all deletion lines compared to the control. For mating phenotypes, we also compared *208-ΔG* females x DGRP_208 males and DGRP_208 females x *208-ΔG* males to the control. We used Fisher’s exact tests to assess the proportion of flies that mated and the proportion of flies that recovered from a chill-induced coma in mating and chill coma recovery time, respectively using the R *stats* package. Models were also run separately by sex. For development time, models were also run separately for *Uhg4*/*CyO* heterozygotes and *Uhg4* homozygotes.

### Ovary Dissection

We placed mated females from representative *Uhg4* deletion lines and their controls (*208-ΔF*, *208-ΔG*, DGRP_208) in fresh food vials supplemented with yeast paste every 12 hours for 36 hours prior to dissection. Flies were dissected in 1X PBS and ovarioles were gently separated. Ovaries were fixed in 4% paraformaldehyde for 15 minutes, followed by three 15-minute washes in PBS with 0.2% Triton X-100. Following a final 15-minute wash in PBS, ovaries were stained with DAPI (Invitrogen) (1μg/mL) for 10 minutes and mounted with ProLong Gold (Invitrogen) immediately after the final PBS wash. Ovaries were imaged on an Olympus Fluoview FV3000 microscope at 20x magnification. Images were processed in Fiji [102].

### RNA Sequencing

We prepared libraries for RNA sequencing from each RNA sample used in the RT-qPCR analyses according to Universal RNA-Seq with NuQuant + UDI (Tecan Genomics, Inc., Redwood City, CA) manufacturer instructions. Specifically, 100ng of total RNA was converted into cDNA via integrated DNase treatment. Second strand cDNA was fragmented using a Covaris ME220 Focused-ultrasonicator (Covaris, Woburn, MA) to 350bp. Drosophila AnyDeplete probes were used to deplete remaining ribosomal RNA and final libraries were amplified using 17 PCR cycles. We used TapeStation High Sensitivity D1000 Screentape (Agilent Technologies, Inc., Santa Clara, CA) and a Qubit™ 1X dsDNA HS Assay kit (Invitrogen, Carlsbad, CA) to measure the final library insert sizes and concentration, respectively. Final libraries were diluted to 4nM and sequenced on a NovaSeq6000 (Illumina, Inc., San Diego, CA) using an S1 flow cell. We sequenced two biological replicates per sex per line (*208-ΔA*, *208-ΔF*, *208-ΔG*, DGRP_208), with ∼20-74 million reads per sample.

Barcoded reads were demultiplexed using the NovaSeq Illumina BaseSpace sequencing pipeline and merged across S1 flow cell lanes. We used the *AfterQC* pipeline (version 0.9.7) [103] to filter out low-quality, short, and adapter reads and the bbduk command within the BBTools package [104] to detect levels of rRNA contamination. We used GMAP-GSNAP [105] to align filtered reads to the *D. melanogaster* reference genome v6.13 and the featurecounts pipeline from the Subread package [106] to count unique alignments to Drosophila genes. Expression for novel transcribed regions (NTRs) was estimated by first compiling a list of NTRs detected from RNA sequencing of young adult DGRP flies [82]. The coordinates of these NTRs were converted from R5 to R6 using the Coordinate Converter tool on FlyBase. A new gene transfer format file was constructed using the coordinate-converted NTR gene models and was used in conjunction with the alignment files for expression estimation. Counts data for each sample were filtered to omit genes for which the median count was less than two, as well as genes for which the proportion of null values across all samples was less than 0.25. The data were then normalized for gene length and library size using Ge-TMM [107]. Filtered and normalized data were analyzed for differential expression using the “PROC glm” command in SAS v3.8 (Cary, NC) according to the ANOVA model *Y = μ* + *L* + *S* + *L*×*S* + *ε*, where Line (*L*), Sex (*S*), and Line×Sex (*L*×*S*) are fixed effects, *Y* is gene expression, *μ* is the overall mean, and *ε* is residual error. A false discovery rate (FDR) correction using the Benjamini-Hochberg procedure (BH-FDR) for multiple tests was applied across all genes to determine the subset of differentially expressed genes significant at BH-FDR < 0.05 and BH-FDR < 0.1 for either the Line or LinexSex terms. NTR expression counts were analyzed using the same approach described above but as a separate dataset. For the bulk RNA sequencing data, plotting ordered raw *p*-values and BH-FDR adjusted *p*-values against the number of tests revealed a non-monotonic relationship between raw *p*-values and adjusted *p*-values. Therefore, we used a BH-FDR thresholding approach to identify genes with statistically significant *p*-values from the ANOVA model. Briefly, after ordering the genes based on ascending raw *p*-values, we compared each gene’s raw *p*-value to FDR critical values calculated as rank*Q/number of tests at both Q = 0.05 and Q = 0.1. For both critical values, *p*-value thresholds were determined as the first occurrence of the raw *p*-values greater than critical values. Genes with raw *p*-values below the *p*-value threshold at critical values Q = 0.05 and Q = 0.1 were considered for downstream analyses. We used the resulting 180 genes significant at FDR < 0.1 for network construction and included the 13 differentially expressed NTRs (BH-FDR < 0.1) with these 180 genes for k-means clustering.

We performed k-means clustering (k = 8, average linkage algorithm) on the Ge-TMM-normalized least squares means of the 193 coregulated genes. We performed iterative k-means clustering with different k values to determine the largest number of clusters without redundant expression patterns across clusters. We used Cytoscape v3.9.1 and the interaction networks from FlyBase [79] (FB_2021_05) to create protein and genetic interaction networks including first-degree neighbors, clustered via MCODE score [108] where applicable. Cluster annotations are based on significantly enriched Gene Ontology (GO) terms. We performed GO statistical overrepresentation analyses with GO: Biological Process Complete, Molecular Function Complete, and Reactome Pathway terms (GO Ontology database released 2021-11-16) using Panther db v16.0 [109] using Fisher Exact tests with BH-FDR correction.

## Supplementary Information Legends

**Table S1. Analyses of behavioral phenotypes.** Analyses of variance (ANOVA) and Fisher’s exact tests comparing *Uhg4* deletion flies to their respective controls.

**Table S2. Differentially expressed genes.** (A) BH-FDR correction for genes based on *Line* and (B) *LinexSex* model terms; (C) Normalized average read counts values for the 180 coregulated genes and 13 NTRs; (D) FDR correction for NTRs genes based on *Line* and *LinexSex* model terms. See corresponding Figure S3.

**Table S3. Analysis of differential expression.** For each gene (A) and NTR (B), the FBgn number or NTR ID, raw *p*-values, BH-FDR adjusted *p*-values for the effects of *Line* (*L*), *Sex* (*S*), and *Line*×*Sex* (*L*×*S*) according to the ANOVA model *Y = μ* + *L* + *S* + *L*×*S* + *ε*.

**Table S4. Gene Ontology enrichment analyses.** Enriched Gene Ontology (GO) Biological Process, Molecular Function and REACTOME terms and their raw and BH-FDR adjusted *p*-values. Groups of genes analyzed include 20 genes/NTRs with differential expression for the *Line* or *Line*×*Sex* terms (BH-FDR < 0.05), 20 genes/NTRs and their 1-degree neighbors, 193 coregulated genes/NTRs (BH-FDR < 0.1) due to a significant *Line* or *Line*×*Sex* effect, 193 genes/NTRs and their 1-degree neighbors, genes within each MCODE cluster (Figure 4, Figure S5), and genes in each k-means cluster (Figure S4).

**Table S5. Gene lists.** Lists of gene names, symbols, and Flybase IDs (FBID) for each network cluster and k-means cluster.

**File S1.** Video of a representative *Uhg4* deletion (*208-ΔG*) female fly displaying an elevated, splayed resting wing position compared to a wildtype female fly. The *Uhg4* deletion fly is walking during the first part of the video and the wildtype fly is walking at the end of the video.

**File S2.** Video of a representative *Uhg4* deletion (*208-ΔG*) male fly displaying an elevated, splayed resting wing position compared to a wildtype male fly. The *Uhg4* deletion fly is to the right of the wildtype fly throughout the video.

**Figure S1. Crossing scheme to generate *Uhg4* deletion CRISPR mutants.** Following injections of *Cas9* and gRNA vectors into embryos of each DGRP line, resulting progeny were screened for presence of a deletion around *Uhg4*, and if a deletion was present, crossed to the original genetic background to generate additional flies heterozygous for the original deletion. Backcrossing siblings heterozygous for the same mutation resulted in homozygous flies that were sterile for all isolated deletion mutations. Virgin female flies heterozygous for a specific deletion were then crossed to male flies containing the *CyO* balancer. Resulting progeny were screened for the presence of the *CyO* balancer chromosome as well as the respective mutation and crossed to full siblings to establish the stock. All females were crossed as virgin flies. “*Δ*” refers to a specific *Uhg4* deletion.

**Figure S2. Mating phenotypes of *Uhg4* deletion flies.** Boxplots displaying time, in seconds, of (A) mating latency (time until mating begins) and (B) mating duration (length of copulation) for each of the *Uhg4* mutant lines (*208-ΔA*, *208-ΔF*, *208-ΔG*), the control line DGRP_208 (*208wt*), and a representative mutant (*208-ΔG*) versus DGRP_208 wildtype pairings (*208-ΔG* females x *208wt* males, *208wt* females x *208-ΔG* males). N = 22-24 pairings of 3-5 day old virgin flies per line. Only flies which successfully initiated or completed mating within 30 minutes were included in analysis for mating latency and mating duration, respectively. See Table S1A. * *p*<0.05

**Figure S3. Non-monotonic relationship between raw and Benjamini-Hochberg adjusted p–values.** (A) raw *p*-values plotted against number of tests; (B) BH-FDR thresholds on raw *p*-values. Raw and adjusted *p*-values are shown in blue and red, respectively. “i”: rank, “m”: number of tests. See corresponding Table S2.

**Figure S4. K-means clusters**. K-means clusters derived from genes and NTRs with differential expression in a global transcriptomic analysis using genes significant for the *Line* or *Line*×*Sex* terms (BH-FDR < 0.1). *Uhg4* is shown in bold and indicated with a star symbol. Magenta indicates a relative higher degree of expression, blue indicates a relative lower degree of expression.

**Figure S5. Additional interaction networks**. Interaction networks containing physical and genetic interactions with genes with a significant *Line* effect. (A) Networks generated from an input of 20 differentially expressed genes/NTRs (BH-FDR < 0.05) including neighbors within at least 1 degree. (B) Networks generated from an input of 180 coregulated genes (BH-FDR < 0.1) including neighbors within at least 2 degrees. Annotation is based on enriched Gene Ontology terms. Dark green indicates the genes in the input data set and light green indicates interaction neighbors. Names are Drosophila gene symbols. See Table S4.

## DATA AVAILABILITY

All high throughput sequencing data are deposited in GEO GSE199865.

Sanger sequencing, qPCR, and raw behavioral data are available on GitHub at rebeccamacpherson/Pleitropic_fitness_effects_Uhg4_rawdata.

## ACKNOWLEDGEMENTS

We thank Miller Barksdale and Patrick Freymuth for their technical assistance, Dr. Sneha Mokashi for experimental advice and Jonathon Thomalla and Drs. Mariana Wolfner, Melissa White, and Deeptiman Chatterjee for their insights into ovary biology. Additionally, we would like to thank the Greenwood Genetic Center and Dr. Heather Flanagan-Steet for the use of their Sanger sequencing and microscope facilities.

## AUTHOR CONTRIBUTIONS

RAM – conceptualization, formal analysis, funding acquisition, investigation, visualization, writing – original draft preparation, writing – review & editing; VS – data curation, formal analysis, software; LS – investigation; RCH – investigation; MRCIII – investigation; RRHA – conceptualization, writing – review & editing, funding acquisition, supervision; TFCM – conceptualization, writing – review & editing, funding acquisition, supervision, project administration.

