## Supplementary figures and images for "Pleiotropic fitness effects of the lncRNA *Uhg4* in *Drosophila melanogaster*"

### Supplemental Figure 1

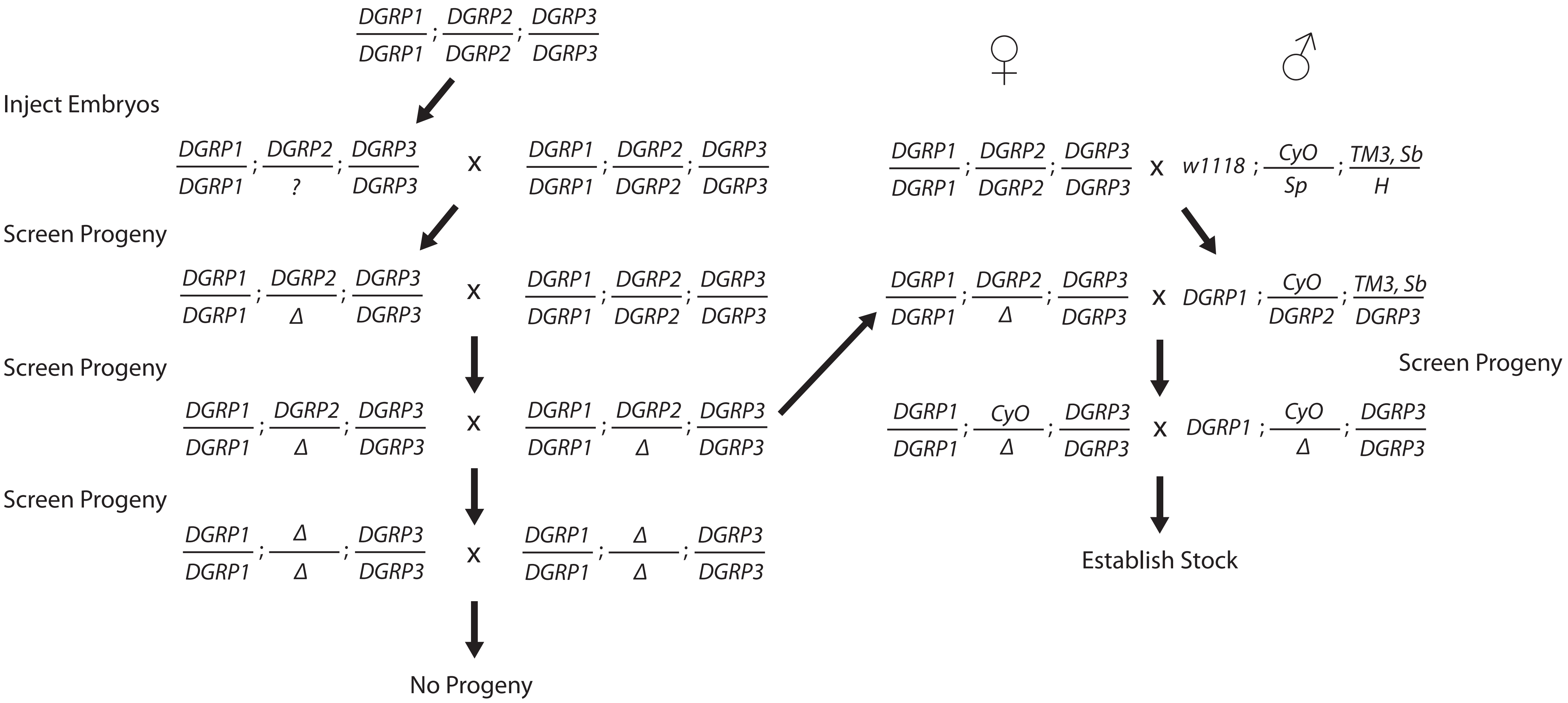

### Supplemental Figure 2

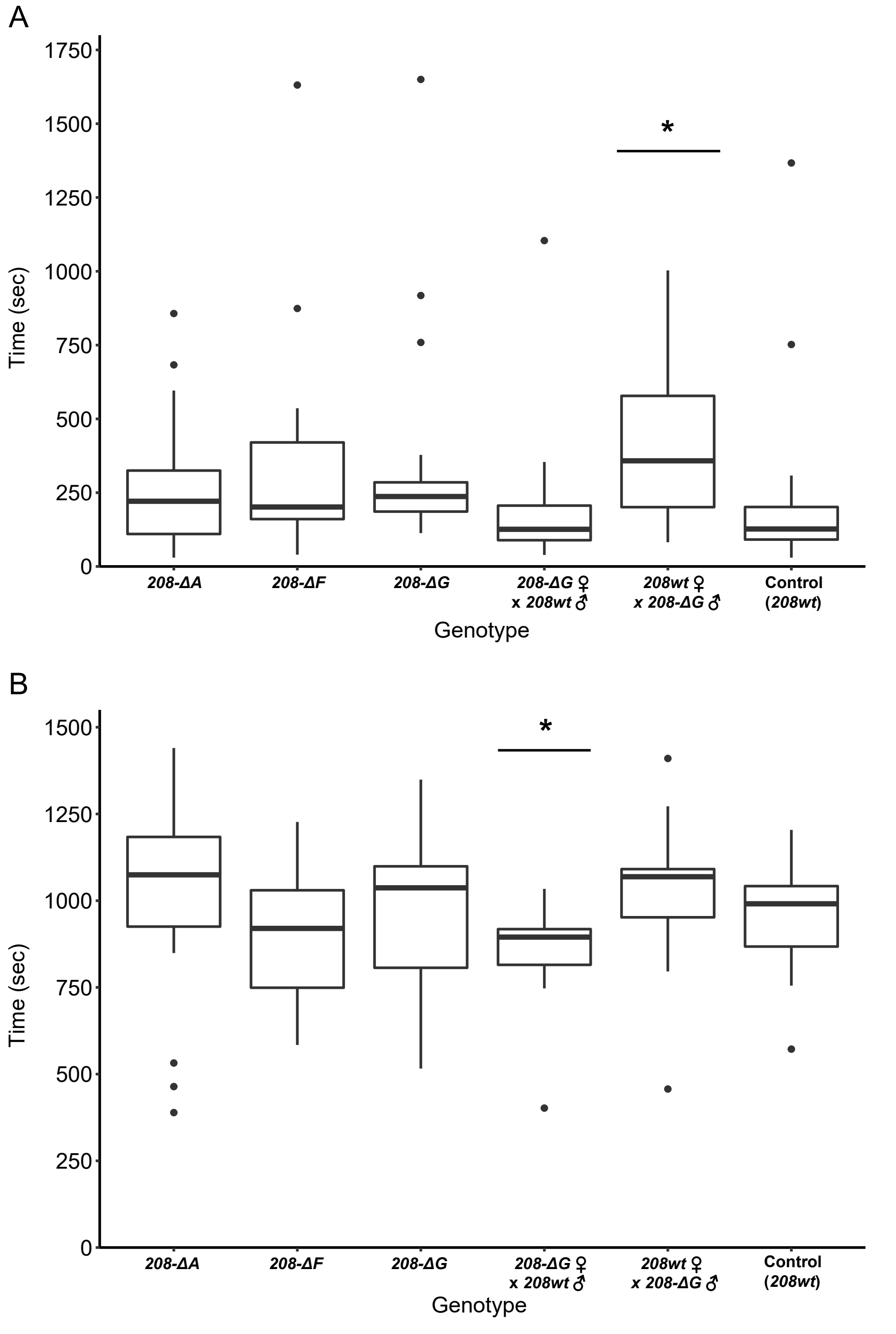

### Supplemental Figure 3

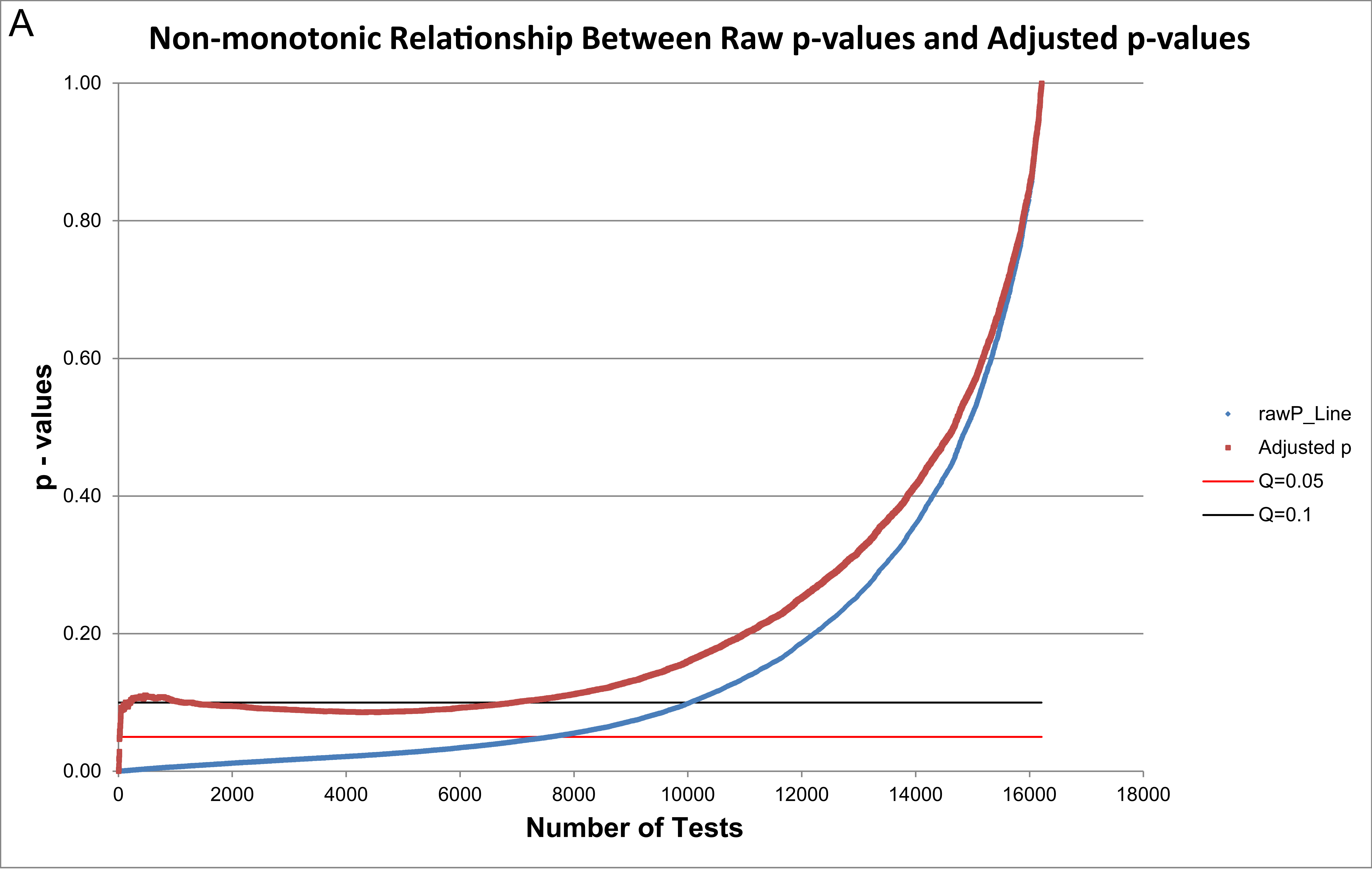

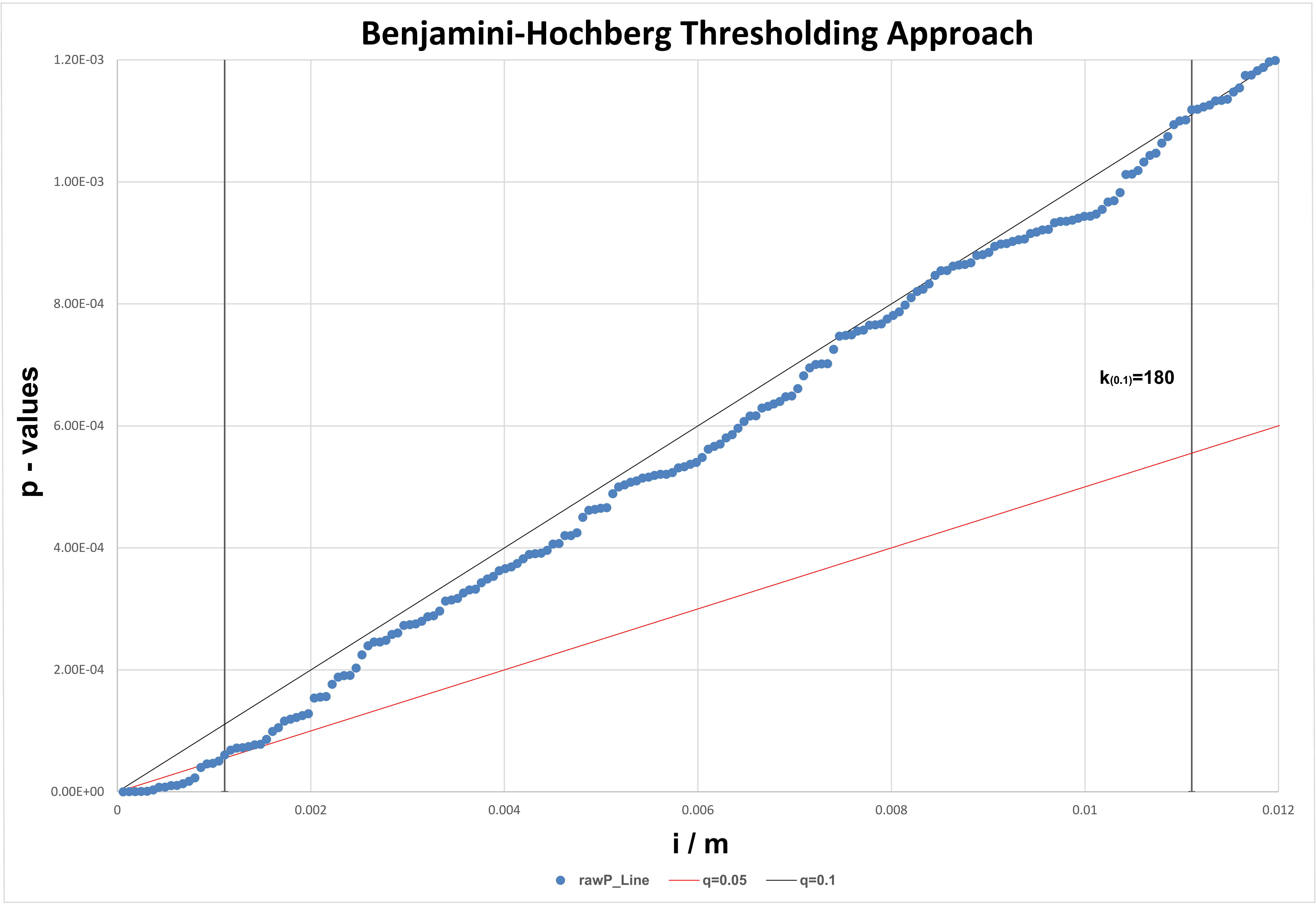

### Supplemental Figure 4

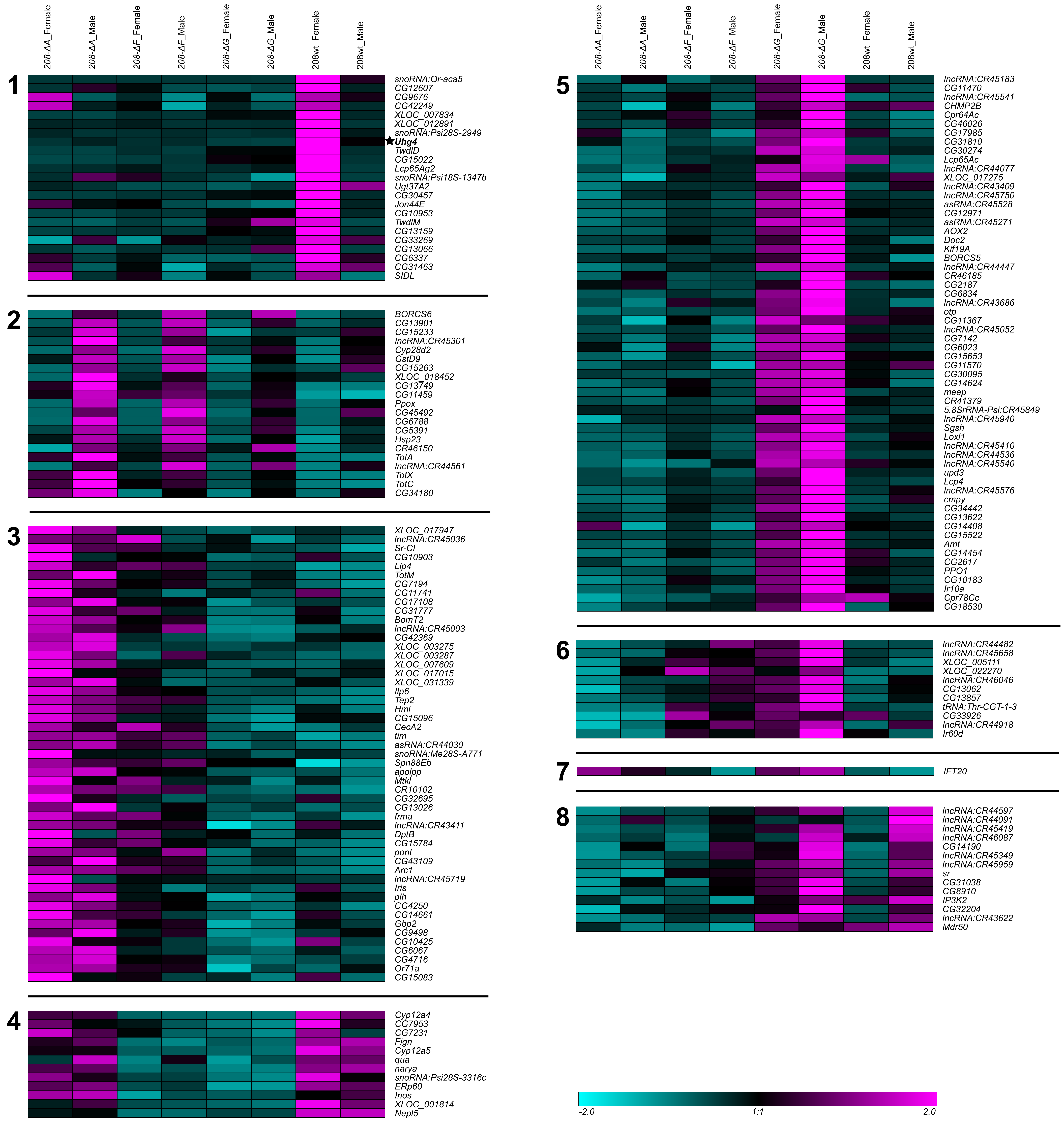

### Supplemental Figure 5

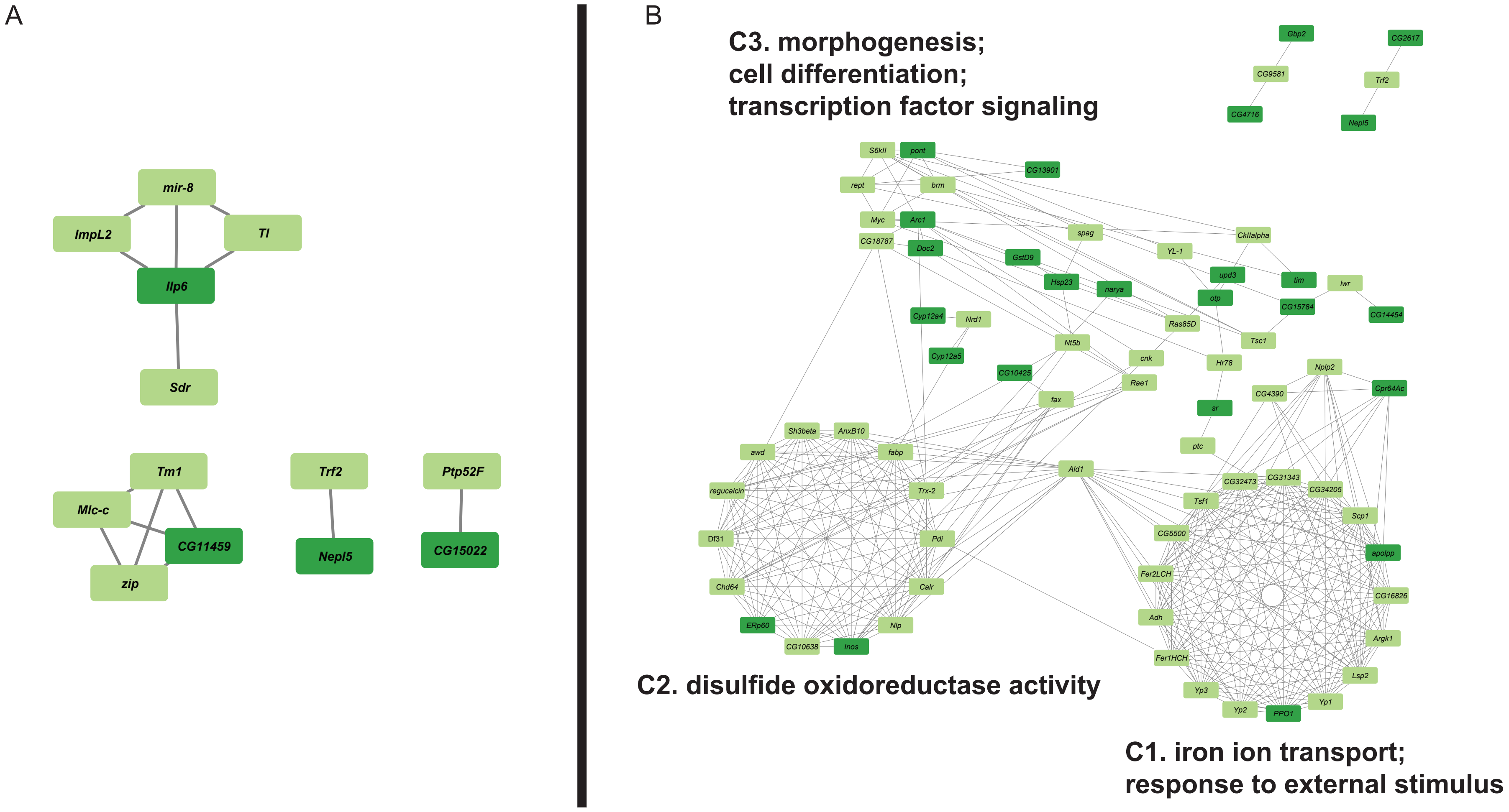
